# Comprehensive Transcriptomic Analysis of Hepatocellular Carcinoma: Uncovering Shared and Unique Molecular Signatures Across Diverse Etiologies

**DOI:** 10.1101/2024.11.23.625031

**Authors:** Babak Khorsand, Nazanin Naderi, Seyedeh Sara Karimian, Maedeh Mohaghegh, Alireza Aghaahmadi, Seyedeh Negin Hadisadegh, Mina Owrang, Hamidreza Houri

**Affiliations:** Department of Neurology, University of California, Irvine, CA, USA; Department of Cell and Molecular Biology, Faculty of Life Science and Biotechnology, Shahid Beheshti University, Tehran, Iran; Department of Oncology, University of Alberta, Canada; Department of Biological Sciences, Islamic Azad University, North Tehran Branch, Tehran, Iran; Department of biology, Islamic Azad university of central Tehran Branch, Tehran, Iran; Department of Chemistry and Chemical Biology, Rutgers, The State University of New Jersey, Piscataway, New Jersey, USA; Faculty of Medical Science, Sari Branch, Islamic Azad University, Sari, Iran; Foodborne and Waterborne Diseases Research Center, Research Institute for Gastroenterology and Liver Diseases, Shahid Beheshti University of Medical Sciences, Tehran, Iran; Celiac Disease and Gluten Related Disorders Research Center, Research Institute for Gastroenterology and Liver Diseases, Shahid Beheshti University of Medical Sciences, Tehran, Iran

**Keywords:** Differentially expressed genes, Hepatocellular carcinoma, Hepatitis B virus, Hepatitis C virus, Liver cirrhosis

## Abstract

Hepatocellular carcinoma (HCC) is a leading cause of cancer mortality, often diagnosed at advanced stages where treatment options are limited. This study undertakes a comprehensive meta-analysis of gene expression profiles from 19 independent datasets sourced from the Gene Expression Omnibus (GEO), encompassing a diverse range of HCC etiologies, including HBV and HCV infections, cirrhosis, and normal liver comparisons. Our analysis identified over 9,000 differentially expressed genes (DEGs), with 125 genes consistently altered across multiple datasets, underscoring their potential as critical biomarkers for HCC. Notably, we observed significant dysregulation in pathways related to cell cycle regulation, immune response, and metabolic processes. The integration of these DEGs across various HCC subtypes provides novel insights into the molecular heterogeneity of HCC, offering promising avenues for the development of targeted therapies and personalized medicine. This extensive repository of DEGs serves as a valuable resource for the scientific community, facilitating further research into the underlying mechanisms of HCC and the pursuit of improved diagnostic and therapeutic strategies.

## 1. Introduction

Hepatocellular carcinoma (HCC) is the most common primary liver cancer, originating from hepatocytes. Globally, HCC is the third most prevalent cancer and the third leading cause of cancer-related deaths [1]. It constitutes over 80% of liver cancer cases, with intrahepatic cholangiocarcinoma (iCCA) and other specified histologies accounting for 14.9% and 5.1% of cases, respectively [2]. HCC development is driven by a complex interplay of environmental and genetic factors. The typical onset of HCC occurs between ages 55 and 65, with variations across different populations, and it disproportionately affects males more than females [3]. Several risk factors are associated with HCC, including liver cirrhosis, chronic infections with hepatitis B virus (HBV) and hepatitis C virus (HCV), excessive alcohol consumption, exposure to aflatoxin B1, and nonalcoholic steatohepatitis (NASH) [4].

Treatment options for HCC are limited to early-stage patients and include tumor resection, liver transplantation, and liver ablation. However, approximately 50% of HCC patients require systemic therapy, such as immunotherapy with immune checkpoint inhibitors (ICIs) and anti-angiogenic monoclonal antibodies like ramucirumab [2, 5, 6]. The prognosis of HCC patients largely depends on the stage at diagnosis. Early detection is associated with improved overall survival, but most patients are diagnosed at advanced stages, precluding curative treatments like hepatic resection and liver transplantation, resulting in a 5-year survival rate of only 18% [7]. Therefore, early detection and timely intervention are crucial for improving survival rates and quality of life for HCC patients. The use of biomarkers for predicting HCC, determining cancer stages, and targeting drug therapies holds significant potential for enhancing survival outcomes [8].

Over the past decade, advancements in tumor biology have been significantly propelled by microarray-based gene expression and RNA sequencing (RNA-seq) analyses, which have shed light on the molecular and genomic mechanisms underlying HCC [9–14]. These studies have identified key genetic and genomic markers of HCC, including the dysregulation of genes involved in cell cycle regulation, the Wnt/β-catenin pathway, cell adhesion molecules critical for cell-cell and cell-matrix interactions, and genes responsible for detoxification and immune response [7, 15–17]. However, it remains unclear whether the integration of these gene dysregulations is also associated with the multiple etiologies of HCC and normal liver. More importantly, the identification of clinically significant transcriptomic changes in HCC across various etiologies reveals substantial variability in patient responses to immunotherapy. This variability complicates the development of effective immunotherapeutic strategies for different HCC subtypes. Single-cell RNA-seq (scRNA-seq) enables a detailed examination of oncogene dysregulations and their interactions at the cellular level. However, despite numerous scRNA-seq studies on HCC, most focus on specific etiologies or a limited number of subtypes. Consequently, the full extent of single-cell heterogeneity in HCC remains largely unexplored.

To address this knowledge gap and elucidate the molecular mechanisms of the identified cell types, this study aims to compile and reanalyze RNA-sequencing and microarray datasets from the Gene Expression Omnibus databases. This comprehensive analysis encompasses 19 studies focused on gene expression profiling in HCC, with the goal of reporting differentially expressed genes (DEGs). The primary objective is to create an extensive repository of data resources that align with the multifaceted dimensions of HCC research, thereby facilitating streamlined data retrieval for researchers engaged in various aspects of HCC studies.

## 2. Material and Methods

### 2.1. Benchmark dataset

This study utilized RNA sequencing and microarray datasets from the Gene Expression Omnibus (GEO) database, encompassing 19 distinct studies focused on gene expression profiling in HCC across various contexts, including early-stage, recurrent, and metastatic disease. The datasets cover a broad spectrum of sample types and clinical scenarios, providing a comprehensive basis for gene expression analysis in HCC.

### 2.2. Summary of Included Datasets

#### 2.2.1. Samsung Medical Center Microarray Data (GSE36376)

Microarray data from 240 patients treated at Samsung Medical Center, Seoul, Korea (July 2000 to May 2006) were analyzed to identify gene expression signatures in HCC post-curative hepatectomy among young patients [18]. All patients had Child-Pugh Class A liver function. The analysis revealed 69 differentially expressed genes, mainly involved in cell cycle and cell division, differentiating patients aged ≤40 years from those >40 years.

#### 2.2.2. RNA-Sequencing Data from Seoul National University Hospital (GSE77509)

This dataset comprises RNA sequencing data from 62 early malignant HCC tumor samples, including 38 patients who underwent surgical resection, alongside 15 normal and 47 adjacent non-tumor samples from patients with HCC or chronic liver disease at Seoul National University Hospital [19]. The study aimed to elucidate transcriptomic profiles during HCC development, revealing a comprehensive immune landscape and dynamic shifts in transcriptomes across 22 major immune cell types from non-tumorous to early malignant stages. Regulatory T cell (Treg) infiltration was noted, with a potential association between CYB561 gene expression and adverse outcomes such as angioinvasion or tumor relapse in patients undergoing total hepatectomy.

#### 2.2.3. HBV-Associated Early-Stage HCC Transcriptome (GSE124535)

This dataset includes 35 HBV-associated early-stage HCC tissues (29 males and 6 females, average age: 55) and 35 healthy tissues collected from Zhongshan Hospital, Fudan University, and Cancer Hospital & Institute, Peking University [20]. The analysis identified eight hub genes (upregulated: AURKB, CDK1, CDC20, CCNB1, CCNA2; downregulated: EHHADH, APOA1, UBB) and key signaling pathways, including metabolic pathways, complement and coagulation cascades, PPAR signaling, and fatty acid degradation. Notably, CDK1, CDC20, CCNB1, and CCNA2 were significantly involved in the cell cycle and p53 signaling pathway.

#### 2.2.4. Genome-Wide Gene Expression in HCC (GSE20140)

Genome-wide gene expression profiles from 287 HCC patients (tumor and adjacent non-tumor cirrhotic tissues) were analyzed, collected from four institutions within the HCC Genomic Consortium: Mount Sinai School of Medicine, New York; IRCCS Istituto Nazionale Tumori, Milan; Hospital Clinic, Barcelona; and Toranomon Hospital, Tokyo [21]. The study examined 22 gene signatures with prognostic value for early-stage HCC, predicting early and overall recurrence in conjunction with clinical and pathological findings.

#### 2.2.5. Pro-Oncogenic Pathways in HCC (GSE1898, GSE4024, GSE9829, GSE14520)

This analysis investigated pro-oncogenic pathways in primary tumors and adjacent non-malignant tissues using genome-wide mRNA expression profiles from 321 HCC patients [22]. Samples included 115 primary tumors and 52 adjacent tissues from the National Cancer Centre of Singapore/Sing Health Tissue Repository, and 206 HCC patients from the Liver Cancer Institute, Fudan University, Shanghai (GSE14520). The study identified 24 ribosomal genes as significant in both tumor and adjacent tissues, with DKK1 serving as a biomarker for poor outcomes. The co-transcriptional signature of ribosome biogenesis genes suggests a novel predictive system.

#### 2.2.6. Genomic Predictors for HCC Recurrence (GSE12720, GSE15239, GSE39791, GSE22058, GSE14520)

This compilation includes microarray data from tumor and matched non-tumor tissues from 72 patients undergoing hepatectomy at Dongsan Medical Center, Keimyung University, Korea (cohort 1) and additional data from Queen Mary Hospital, University of Hong Kong (cohort 2, n=96), and Fudan University, China (cohort 3, n=228) [23]. The study identified a hepatocyte injury regeneration (HIR) signature of 233 unique genes, with four genes (RALGDS, IER3, CEBPD, SLC2A3) significantly associated with HCC recurrence and HBV. Network analysis revealed five upstream regulators: NOTCH1, STAT3, PDX1, TP53, and RELA, highlighting a notable interaction between NOTCH1 and STAT3.

#### 2.2.7. Differential Gene Expression Analysis (GSE89377 and GSE114564)

The microarray dataset GSE89377, comprising 85 HCC tissue samples (24 recurrence and 8 non-The microarray dataset GSE89377, containing 85 HCC tissue samples (24 recurrence and 8 non-recurrence), and the RNA-seq dataset GSE114564, including 53 HCC tissue samples (21 recurrence and 32 non-recurrence), were analyzed to identify novel gene signatures associated with HCC recurrence [24]. A total of 2,385 and 5,927 DEGs were found in the two datasets, respectively, with 981 common DEGs identified via Venn diagram. Gene ontology (GO) analysis linked these genes to biological processes such as the Wnt signaling pathway, angiogenesis, and blood coagulation. Gene set enrichment analysis highlighted candidate genes including CETN2, HMGA1, MPZL1, RACGAP1, and SNRPB as predictive markers for HCC recurrence. Additionally, HMGA1 and RACGAP1 emerged as distinct prognostic indicators.

#### 2.2.8. Transcriptome Sequencing Analysis in HCC from Fujian Provincial Hospital (GSE97214)

RNA-seq data from 63 HCC patients at Fujian Provincial Hospital, China, were analyzed to compare HCC tumors and adjacent non-tumorous tissues [25]. The study identified 943 DEGs, with 690 upregulated and 1,253 downregulated genes involved in pathways such as the cell cycle, DNA replication, p53 signaling, and coagulation cascades. Seven fusion genes were detected, with CRYL1-IFT88 recurring in approximately 9.52% of cases, potentially contributing to HCC progression by reducing the tumor suppressor function of IFT88.

#### 2.2.9. RNA-seq Data from Advanced-Stage HBsAg Positive HCC Patients (GSE128274)

RNA-seq data from 21 advanced-stage HCC patients at the Hospital of the University of Science and Technology of China were analyzed to identify mRNA associations with HCC [26]. All patients were HBsAg positive and had not received anti-tumor treatment prior to surgery. Significant upregulation of mRNAs (ZIC5, C12orf75, C1QL1, TMEM74, GNAZ) and downregulation of others (PZP, FAM65C) were observed in tumor tissues compared to adjacent non-tumor tissues. Differential expression was also noted in lncRNAs, miRNAs, and circRNAs, with volcano plots identifying significant differentials in mRNAs, lncRNAs, miRNAs, and circRNAs, highlighting gene-specific hypo-editing and hyper-editing patterns in HCC.

#### 2.2.10. Serum miRNAs as Biomarkers for HCC Diagnosis (GSE22058)

RNA data from 96 HCC patients and 30 healthy controls from Queen Mary Hospital were analyzed to explore circulating miRNAs as biomarkers for HCC diagnosis [27]. The study sought to address the limitations of alpha-fetoprotein (AFP) in HCC diagnosis. Validation with 116 serum samples from Chang Zheng Hospital and Eastern Hepatobiliary Surgery Hospital confirmed the significant reduction of miR-15b and miR-130b post-surgery, indicating their potential as serum biomarkers for HCC diagnosis.

#### 2.2.11. IsomiR-21-5p in Liver Cancer Progression (GSE174608 and GSE114564)

miRNA and RNA sequencing data from 62 patients at Catholic University Hospital and 15 healthy individuals, along with data from TCGA_LIHC and GEO, identified isomiR-21-5p as a potent pro-tumorigenic isomiR in liver cancer development [28].

#### 2.2.12. Lipid Metabolism Reprogramming in HCC (GSE140463 and GSE140243)

RNA-seq data from seven patients with HCC undergoing liver transplantation at Addenbrooke’s Hospital were analyzed to study lipid metabolism reprogramming in cirrhotic fatty liver [29]. An integrated systems biology approach revealed a positive correlation between MUFA-PC levels and genes involved in lipogenesis and PC synthesis, highlighting the significance of MUFA-PC in HCC proliferation.

#### 2.2.13. Microarray Data of Carbon Metabolism in HCC (GSE41804, GSE17548, GSE29721, GSE33006, GSE40873, GSE6222)

Two distinct microarray datasets were utilized to explore gene expression patterns in HCC and non-tumorous liver tissues [30]. Dataset Set-1 (E-MTAB-950) consisted of 120 HCC samples and 160 non-tumor samples, while Set-2 (GSE41804, GSE17548, GSE29721, GSE33006, GSE40873, GSE6222) included 60 HCC samples and 104 non-tumor samples. Additionally, RNA-seq data from three HCV-infected HCC patients (GSE81550) were analyzed to identify a 22-gene signature specifically associated with glycolysis regulation in HCC.

#### 2.2.14. Metabolism-Related Genes in HCC Carcinogenesis (GSE77314)

RNA sequencing data from 50 HCC patients, encompassing both tumor and peri-tumoral tissues, were examined to identify DEGs across different metastasis stages [31]. The analysis revealed 8,900 DEGs in early metastasis stages and 1,789 DEGs in advanced metastasis stages. Notably, the gene DMGDH was found to be downregulated, suggesting its potential role as a predictive marker for HCC and its involvement in inhibiting metastasis via the Akt signaling pathway.

#### 2.2.15. DNA Methylation Patterns in HCV-Associated HCC (GSE82178)

RNA-seq data from paired tumor (n=9) and non-tumor tissues from HCV-infected (n=10) and uninfected individuals (n=11) were analyzed to identify distinct DNA methylation patterns associated with HCC development [32]. The study highlighted the addition of DNA methylation targeted to candidate enhancers in liver cells, particularly enriched at the binding sites of FOXA1, FOXA2, and HNF4A transcription factors. These methylation patterns were implicated in the regulation of genes related to liver cancer and stem cell biology.

#### 2.2.16. RNA Sequencing Data Indicated Prognostic Subtypes in HCC (GSE87630)

In a comprehensive multi-omics analysis, RNA-seq data identified three prognostic subtypes of HCC, designated as iCl1, iCl2, and iCl3, with the iCl1 subtype exhibiting the poorest prognosis [33]. The study linked deviations in CNVcor and METcor genes to unfavorable clinical outcomes, with CA9 emerging as a key differentially expressed gene in more invasive tumor phenotypes.

#### 2.2.17. RNA Sequencing Data of Ribosome Profiling in HCC (GSE112705)

Ribosome profiling data from 10 HCC patients were analyzed, revealing dysregulated translation processes in HCC at sub-codon resolution [34]. The study identified significant variations in translation efficiency and frame usage, which may have important implications for understanding the molecular mechanisms underpinning tumor biology.

#### 2.2.18. Comprehensive Transcriptome Profiling in HCC (GSE105130)

RNA sequencing data from 25 HCC patients, predominantly HBV-positive, were analyzed to identify 53,224 transcripts with significant differential expression between tumor and non-tumor tissues [35]. Pathway analysis indicated enrichment in cell cycle regulation, apoptosis, and DNA repair pathways, with key regulators such as CDK1 and TP73 playing crucial roles in HCC progression.

#### 2.1.19. Oncogenic Markers in HCC Subgroups (GSE14520, GSE1898, and GSE4024)

Gene expression data from 380 HCC patients identified the oncoprotein YY1AP1 as a critical factor in the EpCAM+ AFP+ HCC subtype [36]. YY1AP1’s involvement in chromatin topology and stem cell regulation underscores its potential as a therapeutic target in this aggressive HCC subgroup.

### 2.3. Analysis

DEG analyses were conducted on RNA-seq and microarray datasets, identifying genes with significant upregulation (≥ twofold change, p-value < 0.05) and downregulation (≤ twofold change, p-value < 0.05) in HCC tissues compared to non-HCC controls. Enrichment analyses were performed on the DEGs that reached from the previous step by DAVID [37] to extract enriched DEGs. Finally, to further investigate the molecular interactions underlying HCC and to elucidate the potential mechanisms distinguishing HCC from other liver conditions, we constructed a protein-protein interaction (PPI) network from the enriched DEGs of the previous step.

## 3. Results

To uncover the common DEGs across a range of HCC-related conditions, we conducted an in-depth analysis across 19 distinct dataset groups. These groups encompassed comparisons between HCC and various liver conditions, including HCC HBV-infected tissue versus non-cancerous HBV-infected tissue from seven datasets (GSE112705, GSE14520, GSE15239, GSE22058, GSE77509, GSE87630, and GSE97214); HCC tissue versus normal liver tissue from five datasets (GSE89377, GSE128274, GSE1898, GSE33294, and GSE39791); HCC HCV-infected tissue versus non-cancerous HCV-infected tissue from four datasets (GSE29721, GSE33006, GSE41804, and GSE81550); HCC tissue versus non-cancerous cirrhotic tissue from three datasets (GSE20140, GSE114564, and GSE89377); HCC tissue versus HBV-related cirrhosis from one dataset (GSE17548); HCC tissue versus HCV-related cirrhosis from one dataset (GSE17548); HCC tissue versus chronic hepatitis of various etiologies from one dataset (GSE17548); HCC HBV-infected tissue versus HBV-related cirrhosis from one dataset (GSE17548); and HCC HCV-infected tissue versus HCV-related cirrhosis from one dataset (GSE17548). This comprehensive analysis enabled the identification of DEGs that are consistently altered across multiple HCC conditions, offering valuable insights into the molecular mechanisms underlying HCC. The identified DEGs across these HCC conditions are depicted in a heatmap (Figure 1).

**Figure 1.**
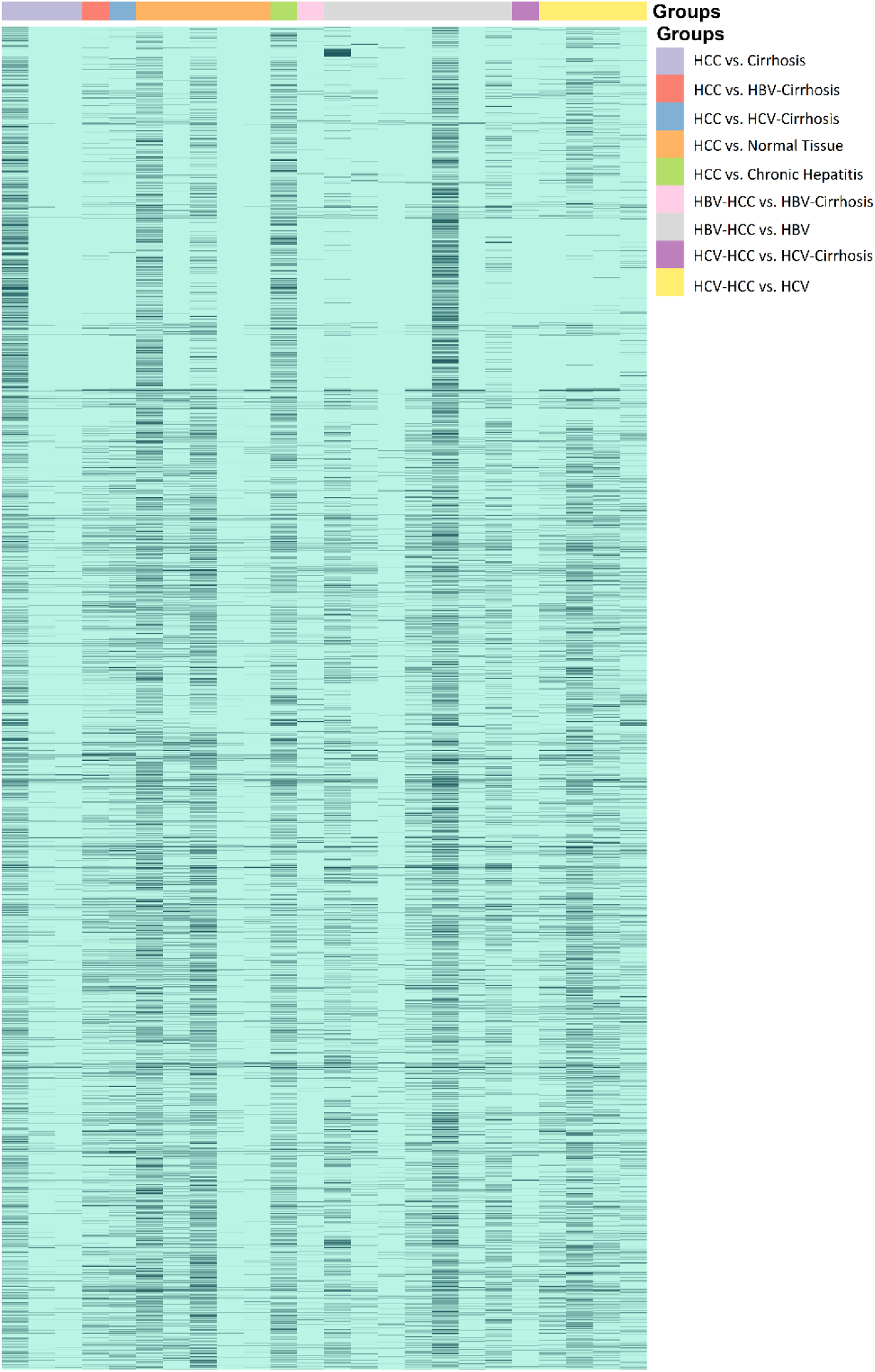
Heatmap illustrating the identified DEGs of HCC (Tumor) compared to various liver conditions.

### 3.1. Differential Gene Expression Analysis Between HCC Tissue and Normal Liver Tissue

To uncover the DEG profiles distinguishing HCC from normal liver tissue, we analyzed data from five independent datasets. Specifically, we identified 627 DEGs in GSE89377, 5,543 DEGs in GSE128274, 2,210 DEGs in GSE1898, 4,923 DEGs in GSE33294, and 477 DEGs in GSE39791. Altogether, this analysis led to the discovery of 9,119 DEGs, which included 4870 genes that were upregulated and 4249 genes that were downregulated, using the criteria of log2 fold change (Log2 FC) > or < 1 and an adjusted *p*-value < 0.05. Comprehensive details of the identified DEGs, along with their expression patterns across these datasets, are provided in Supplementary File 1. Among these, 125 genes consistently exhibited differential expression across all five datasets, indicating their potential as critical biomarkers for differentiating HCC from normal liver tissue. The full list of these common DEGs is presented in Supplementary File 2. Moreover, Figure 2 provides a Venn diagram that visually represents the overlap of DEGs among the datasets comparing HCC tissue with normal liver tissue.

**Figure 2.**
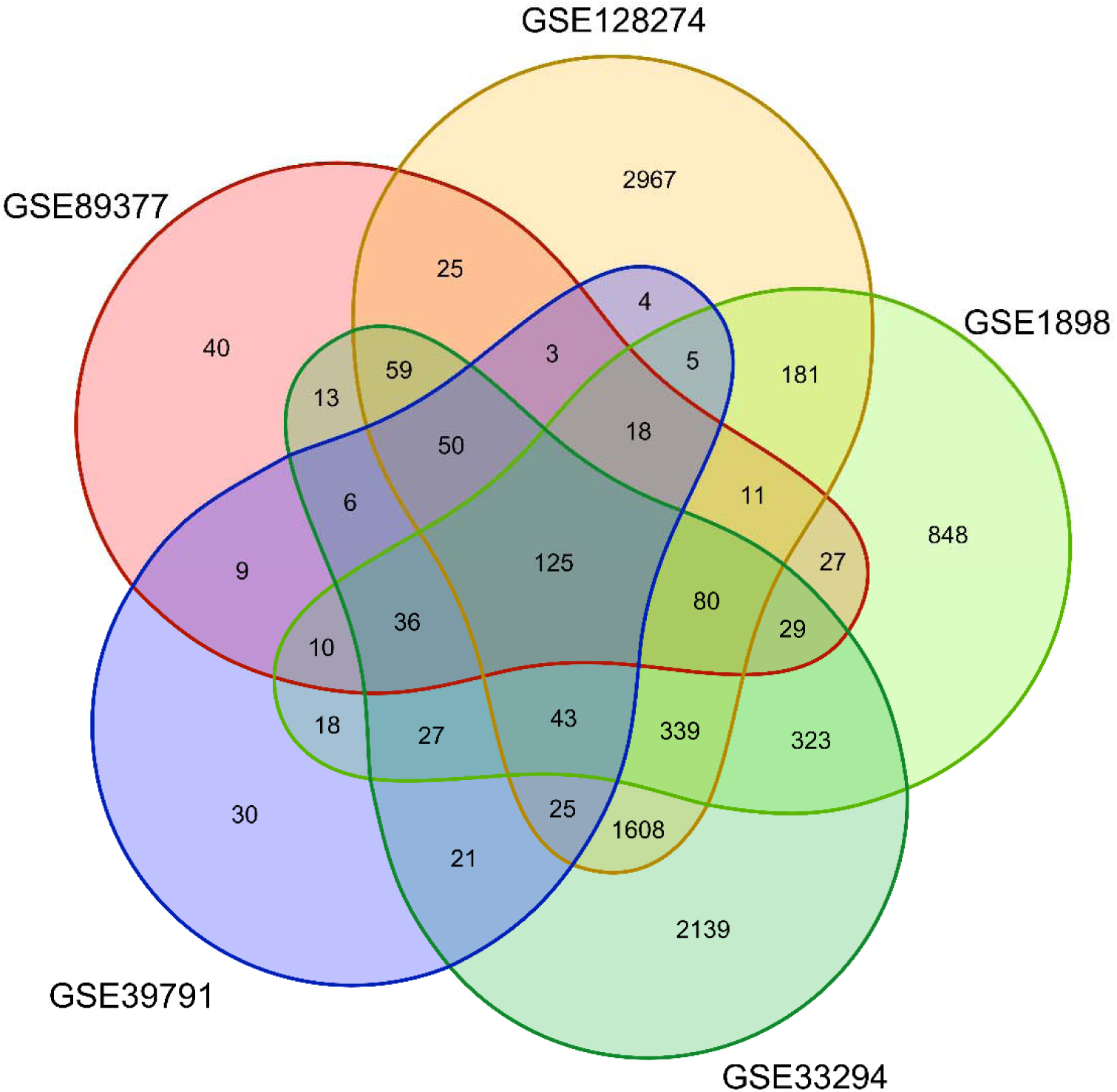
Venn diagram that visually represents the overlap of DEGs among five datasets comparing HCC tissue with normal liver tissue.

### 3.2. Analysis of DEGs in HCC HBV-Infected Tissue Compared to Non-Cancerous HBV-Infected Tissue

To identify DEGs between HCC in HBV-infected patients and their non-cancerous counterparts, we analyze 2,860 DEGs in GSE112705, 1,672 DEGs in GSE14520, 608 DEGs in GSE15239, 2,408 DEGs in GSE22058, 6,784 DEGs in GSE77509, 1,164 DEGs in GSE87630, and 2,804 DEGs in GSE97214. This comprehensive approach led to the identification of a total of 10,018 DEGs, encompassing 6280 upregulated genes and 3738 downregulated genes. A detailed analysis of the expression profiles and the distribution of these DEGs across the datasets is provided in Supplementary File 3. Notably, 14 genes were consistently differentially expressed across all seven datasets, indicating their potential as robust biomarkers for distinguishing HCC in HBV-infected patients from non-cancerous HBV-infected tissue (Supplementary File 2). These key genes include ATP-binding cassette subfamily A member 8 (ABCA8), catalase (CAT), C-C motif chemokine ligand 20 (CCL20), F11, GABA type A receptor-associated protein-like 1 (GABARAPL1), growth arrest and DNA damage-inducible beta (GADD45B), GTP cyclohydrolase 1 (GCH1), metallothionein 1G (MT1G), metallothionein 1M (MT1M), regulator of calcineurin 1 (RCAN1), Rho family GTPase 3 (RND3), S100 calcium-binding protein P (S100P), suppressor of cytokine signaling 2 (SOCS2), and ZFP36 ring finger protein (ZFP36). Additionally, 182 and 288 DEGs were found to be common across six and five datasets, respectively. The Venn diagram in Figure 3 visualizes the intersection of DEGs among the seven datasets, highlighting the genes consistently associated with HCC in HBV-infected patients.

**Figure 3.**
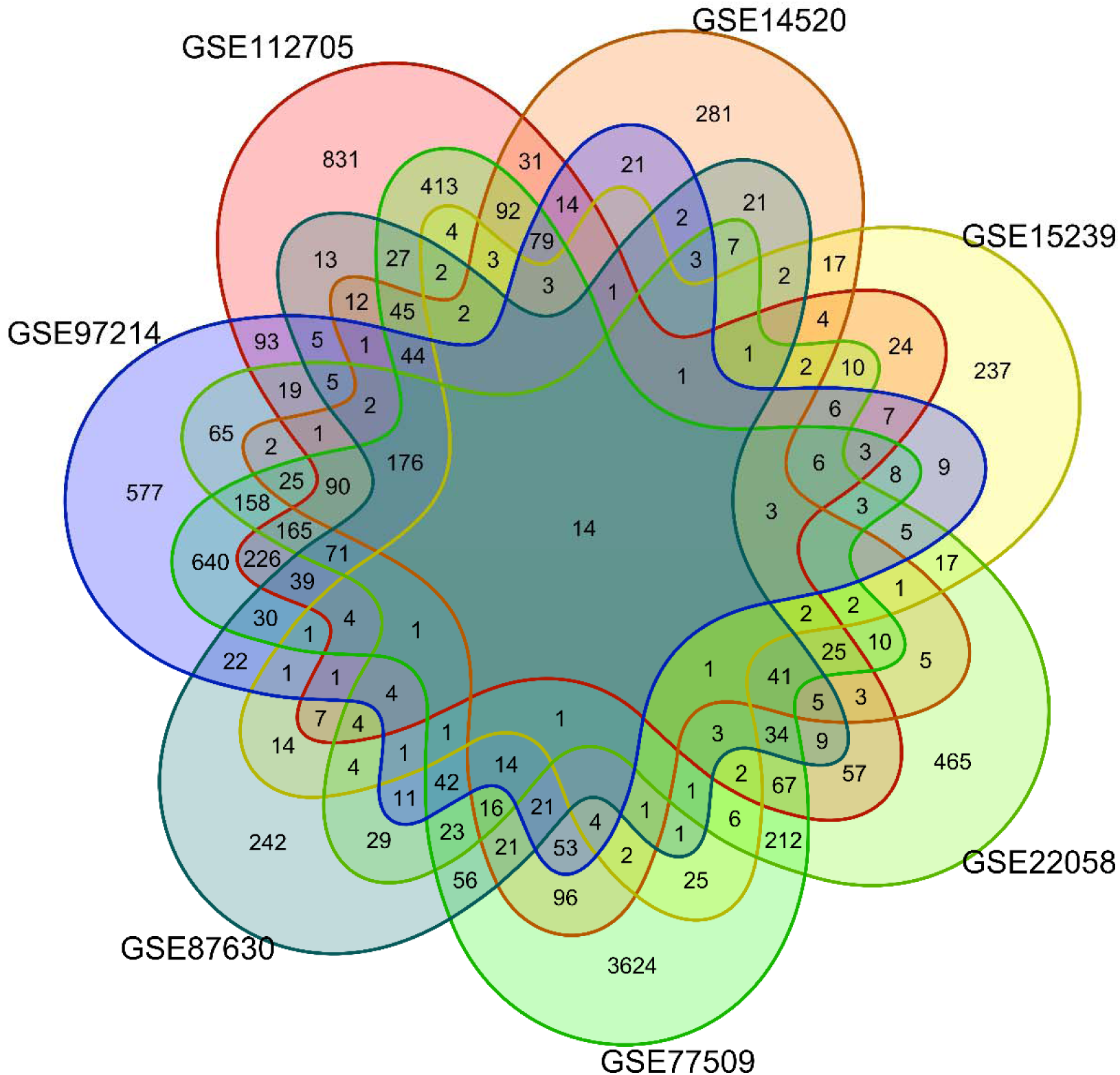
Venn diagram visualizing the intersection of DEGs among the seven datasets, highlighting the genes consistently associated with HCC in HBV-infected patients compared to HBV-infected patients without HCC.

### 3.3. Analysis of DEGs in HCC HCV-Infected Tissue Compared to Non-Cancerous HCV-Infected Tissue

To identify the DEGs between HCC in HCV-infected patients and their non-cancerous counterparts, we conducted an analysis of the following datasets: GSE29721 (2,403 DEGs), GSE33006 (5,309 DEGs), GSE41804 (3,252 DEGs), and GSE81550 (1,790 DEGs). Our comprehensive examination resulted in the identification of 7,403 DEGs in total, comprising 3395 upregulated and 4008 downregulated genes. The detailed expression profiles and distribution of these DEGs across the datasets analyzing HCC in HCV-infected patients and their non-cancerous counterparts are available in Supplementary File 4. Importantly, 221 genes were consistently differentially expressed across all four datasets, underscoring their potential as reliable biomarkers for distinguishing HCC in HCV-infected patients from non-cancerous HCV-infected tissue (as listed in Supplementary File 4). Figure 4 illustrates the Venn diagram depicting the intersection of DEGs among the seven datasets, highlighting the genes consistently associated with HCC in HCV-infected patients.

**Figure 4.**
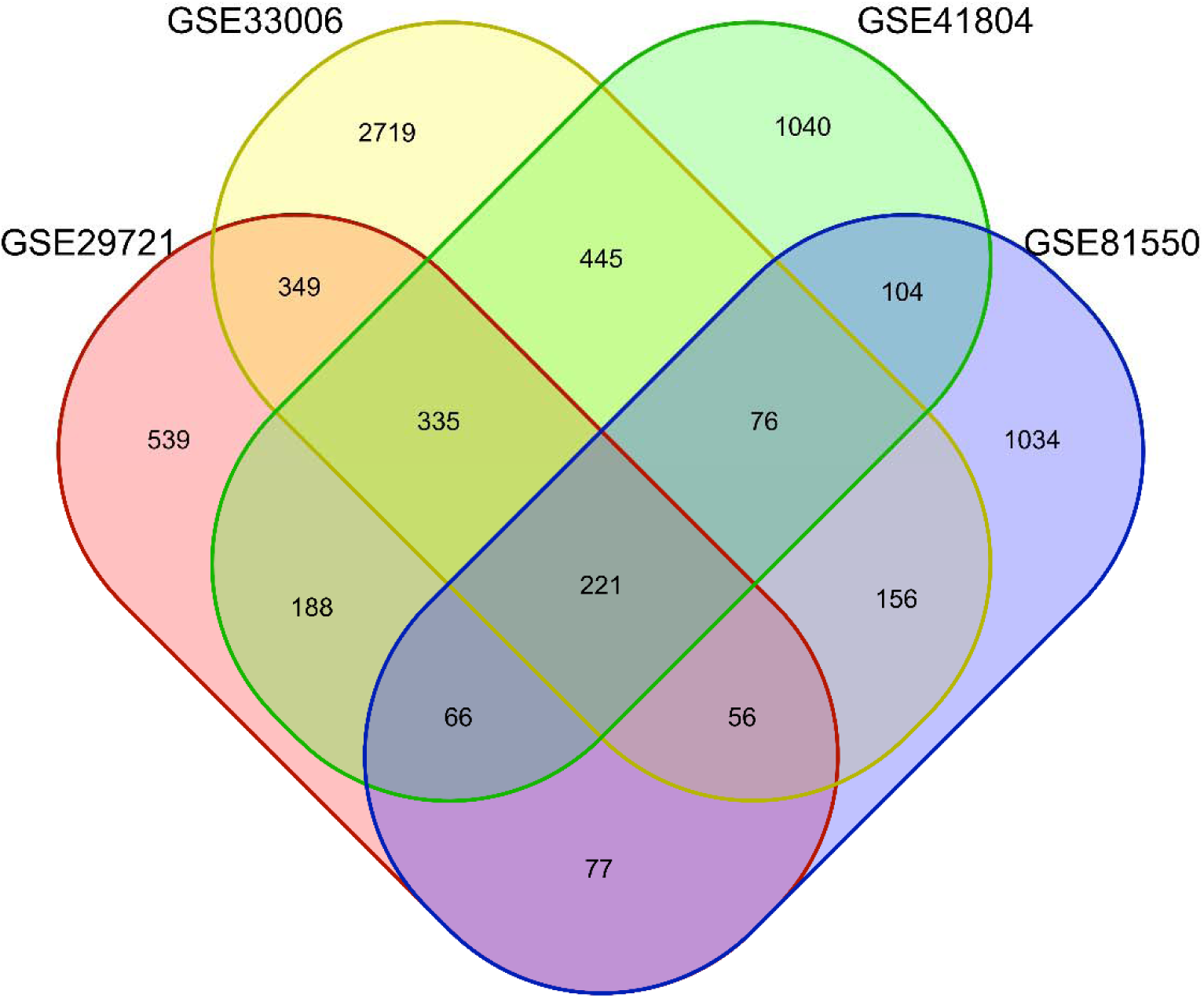
Venn diagram visualizing the intersection of DEGs among the seven datasets, highlighting the genes consistently associated with HCC in HCV-infected patients compared to HCV-infected patients without HCC.

### 3.4. Differential Gene Expression Analysis Between HCC and Cirrhosis

To distinguish DEGs between HCC and cirrhosis, we analyzed data from the following datasets: GSE20140 (648 DEGs), GSE114564 (4,882 DEGs), and GSE89377 (431 DEGs). This comprehensive analysis identified a total of 5,207 DEGs, including 3087 upregulated and 2120 downregulated genes. The detailed expression profiles and distribution of these DEGs across the datasets differentiating HCC from cirrhosis can be found in Supplementary File 5. Notably, 164 genes were consistently differentially expressed across all three datasets, highlighting their potential as robust biomarkers for distinguishing HCC from cirrhosis (as detailed in Supplementary File 5). Figure 5 presents a Venn diagram illustrating the overlap of DEGs among these datasets, emphasizing the genes consistently associated with HCC in comparison to cirrhosis.

**Figure 5.**
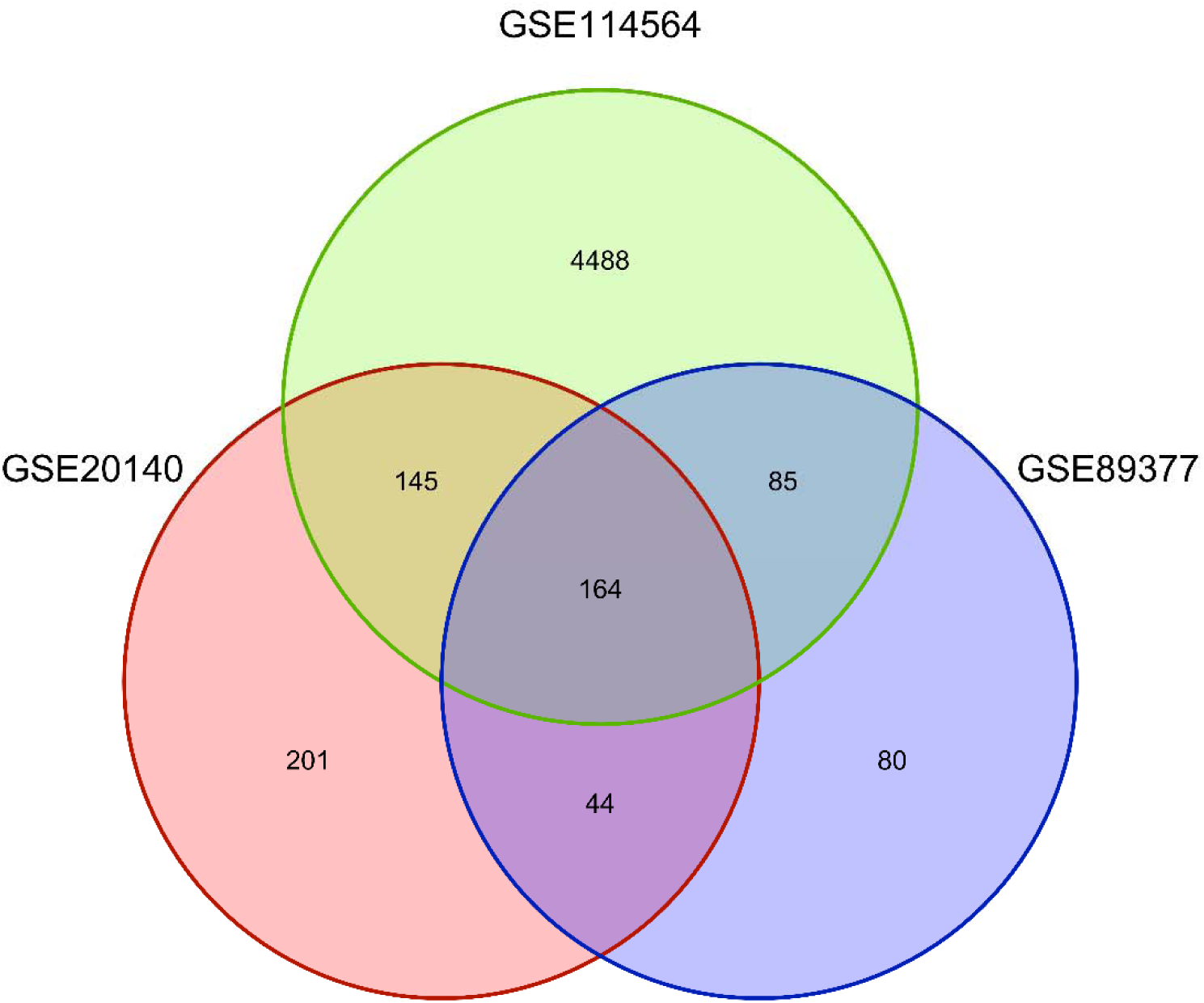
Venn diagram illustrating the overlap of DEGs among three datasets, emphasizing the genes consistently associated with HCC in comparison to cirrhosis

### 3.5. Analysis of DEGs in HCC Compared to Other Liver Conditions

We identified DEGs that are consistently expressed differentially in HCC tissue compared to other liver conditions, including 2,993 DEGs (1393 upregulated vs. 1600 downregulated) distinguishing HCC from HBV-related cirrhosis, 3,568 DEGs (1695 upregulated vs. 1873 downregulated) distinguishing HCC from HCV-related cirrhosis, and 4,403 DEGs (2539 upregulated vs. 1864 downregulated) distinguishing HCC from chronic hepatitis of various etiologies. Furthermore, we identified 939 DEGs (232 upregulated vs. 707 downregulated) differentiating HBV-related HCC from HBV-related cirrhosis. Comprehensive details of these DEGs, including their expression profiles across the analyzed datasets, are provided in Supplementary File 6.

### 3.6. Protein-Protein Interaction Network Construction

The constructed PPI of 64 enriched DEGS, were identified in our comparative analysis. The resulting network, as illustrated in Figure 6, highlights the complex interplay among various proteins encoded by DEGs, revealing key hubs and interaction clusters that may play critical roles in HCC pathogenesis.

**Figure 6.**
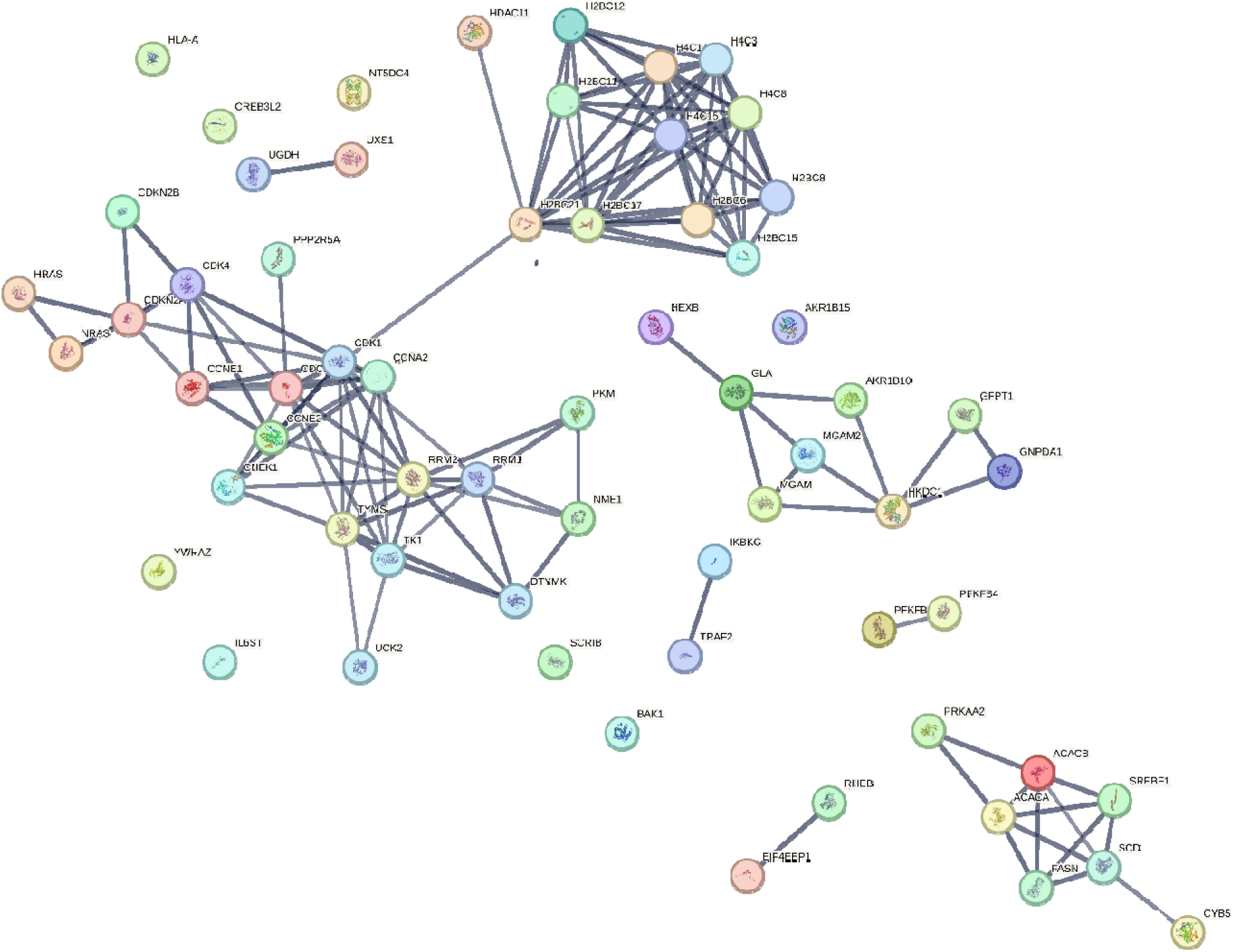
PPI network illustrating interplay among various proteins encoded by DEGs, revealing key hubs and interaction clusters that may play critical roles in HCC pathogenesis.

The PPI network analysis revealed several prominent clusters of interacting proteins, indicating potential pathways and biological processes that are significantly altered in HCC. Notably, several central nodes, including cyclin-dependent kinase 1 (CDK1), cyclin E1 (CCNE1), and thymidylate synthase (TYMS), were identified as major hubs within the network. These proteins are known to be involved in cell cycle regulation, DNA replication, and repair, processes that are frequently dysregulated in cancer, suggesting their pivotal role in HCC development. Furthermore, the interaction network displayed extensive connectivity among proteins related to metabolic pathways, particularly those involved in glycolysis and lipid metabolism. Key enzymes such as hexokinase 2 (HK2), pyruvate kinase M1/2 (PKM), and acetyl-CoA carboxylase alpha (ACACA) were highlighted, reflecting the metabolic reprogramming often observed in cancer cells, known as the Warburg effect. This metabolic shift may contribute to the aggressive nature of HCC and its resistance to conventional therapies. Interestingly, the network also emphasized interactions involving proteins associated with cellular stress responses and immune modulation, such as catalase (CAT), superoxide dismutase 1 (SOD1), and interleukin-6 receptor (IL6R). The dysregulation of these proteins could be linked to the inflammatory microenvironment commonly observed in HCC, especially in the context of underlying liver conditions like cirrhosis and hepatitis. PPI network analysis of DEGs in HCC compared to other liver conditions has uncovered critical molecular interactions that could serve as potential biomarkers or therapeutic targets. The identified hubs and clusters within the network offer insights into the underlying mechanisms of HCC pathogenesis and highlight the complexity of its molecular landscape. Further investigation into these interactions and their functional implications may provide valuable avenues for the development of targeted therapies and improve the clinical management of HCC.

## 4. Discussion

This study undertook a comprehensive multi-database analysis to identify crucial DEGs associated with HCC that differentiate it not only from normal liver tissue but also from other liver conditions such as cirrhosis and viral hepatitis. Our findings suggest that these DEGs have strong potential to serve as independent prognostic biomarkers for HCC patients. By integrating data from 19 distinct dataset groups, encompassing various comparisons between HCC and other liver conditions, we were able to identify a robust set of DEGs consistently distinguishing HCC from both normal and pathological liver tissues. These DEGs are thoroughly documented in Supplementary Files 1-6, providing a valuable resource for researchers seeking to identify potential biomarkers for further experimental validation.

The DEG analysis comparing HCC tissues to normal liver tissues across five datasets highlighted a substantial number of DEGs, reflecting the extensive molecular alterations that occur in HCC. Among these, 125 genes were consistently differentially expressed across all datasets, suggesting their utility as universal biomarkers for HCC. These genes play critical roles in various fundamental cellular processes, including cell cycle regulation and mitosis; signal transduction and cellular communication; metabolic pathways such as amino acid metabolism, fatty acid oxidation, glycolysis, and drug metabolism; ion, nutrient, and metabolite transport across cellular membranes; and cellular stress responses, including oxidative stress and damage repair. Particularly, genes related to retinol and retinoid metabolic processes, such as CYP2C9, CYP2E1, CYP39A1, CYP3A4, CYP4A11, CYP4F2, CYP8B1, RDH5, and RDH16, were significantly enriched among the common DEGs. Previous studies have linked altered retinol homeostasis to various liver diseases, including non-alcoholic fatty liver disease (NAFLD) and NASH, which can progress to hepatic fibrosis and eventually HCC [38, 39]. Additionally, members of the solute-carrier (SLC) gene superfamily, including SLC22A1, SLC27A5, SLC38A4, and SLCO1B3, were identified as important common DEGs. The SLC gene superfamily has been implicated in tumorigenesis, metastasis, and chemoresistance in HCC, suggesting that the dysregulation of these genes may offer novel strategies for HCC diagnosis [40, 41]. Moreover, SLC genes have demonstrated promising accuracy and generalizability in assessing prognosis and predicting survival outcomes for HCC patients[40].

Despite the abundance of research on HCC biomarkers, particularly in the context of HBV-related HCC, there remains a gap in the comprehensive exploration of prognostic biomarkers for this subtype of the disease. Our analysis comparing HCC HBV-infected tissues with non-cancerous HBV-infected liver tissues identified 14 genes consistently differentially expressed across all seven datasets, highlighting their potential as specific biomarkers for HCC in the context of HBV infection. These genes include ABCA8, CAT, CCL20, F11, GABARAPL1, GADD45B, GCH1, MT1G, MT1M, RCAN1, RND3, S100P, SOCS2, and ZFP36. Particularly, ABCA8 and GADD45B emerged as prominent candidates, indicating their significant role in distinguishing HCC from non-cancerous HBV-infected tissues. Other studies have identified prognostic-related genes in HBV-associated HCC, such as TYMS, MAD2L1, CCNA2, CDK1, and SPP1, with some of these genes also emerging as common DEGs in our analysis [42]. Moreover, research by Zeng et al. identified KIF11, TPX2, KIF20A, and CCNB2 as potential independent prognostic genes and diagnostic targets for HBV-related HCC [43]. These findings suggest that the development and progression of HBV-related HCC are closely linked to pathways involving cell cycle regulation, mitosis, p53 signaling, retinol metabolism, and organic acid catabolism.

Similarly, the comparison of HCC HCV-infected tissues with non-cancerous HCV-infected liver tissues revealed 221 DEGs consistently differentially expressed across all four datasets, underscoring their robustness as biomarkers for HCC in HCV-infected patients. Previous studies, such as those by Zhang et al., have reported significant upregulation of cell cycle-related genes in HCV-related HCC, including CDK1, CCNB1, CDC20, NEK2, AURKA, RACGAP1, CDKN2A, CDKN2B, CDKN3, RRM2, and ASPM [44]. Liu et al. further identified 368 DEGs and 10 hub genes as potential diagnostic biomarkers and therapeutic targets for HCV-related HCC, highlighting CCNB1, KIF20A, and HMMR as candidate targets for diagnosis and therapy [45].

The identification of consistent DEGs and key protein interactions offers promising avenues for developing novel biomarkers and therapeutic targets for HCC. The robustness of certain genes and proteins across multiple datasets emphasizes their potential as diagnostic and prognostic tools in HCC. Moreover, understanding the molecular interactions and pathways involved in HCC could pave the way for targeted therapies aimed at disrupting specific pathways altered in this malignancy. Further functional validation of these biomarkers and therapeutic targets is essential to translate these findings into clinical practice and improve patient outcomes.

## 5. Conclusion

our comprehensive DEG analysis and PPI network construction provide a detailed molecular landscape of HCC, offering valuable insights into its pathogenesis and potential therapeutic intervention points. These findings will inform researchers and clinicians in identifying potential biomarkers for the diagnosis and prognosis of HCC, distinguishing it from other liver conditions, and evaluating patient outcomes from chronic hepatitis liver fibrosis or cirrhosis. Future research should focus on validating these findings in clinical settings and exploring the functional roles of the identified genes and proteins in HCC progression and treatment.

## Author contributions

H.H. and B.K. conceptualized the study. M.M., A.A., and S.N.H. conducted the systematic search and data collection. B.K., N.N., S.S.K., and M.M. analyzed the data. M.O. drafted the manuscript, while H.H. supervised the project, reviewed the manuscript, and provided final approval.

## Ethics approval and consent to participate

This study was reviewed and approved by the Institutional Ethical Review Committee of the Research Institute for Gastroenterology and Liver Diseases at Shahid Beheshti University of Medical Sciences (No. IR.SBMU.RETECH.REC.1402.645).

## Consent for publication

Not applicable.

## Availability of data and materials

The data that support the findings of this study are available in the article and related supplementary files.

## Acknowledgments

We are grateful to the members of the Research Institute for Gastroenterology and Liver Diseases, Shahid Beheshti University of Medical Sciences, Tehran, Iran. The authors also really appreciate Professor Mohammad Reza Zali and Dr. Amir Sadeghi for their support.

